# microbeSEG: A deep learning software tool with OMERO data management for efficient and accurate cell segmentation

**DOI:** 10.1101/2022.04.29.489998

**Authors:** Tim Scherr, Johannes Seiffarth, Bastian Wollenhaupt, Oliver Neumann, Marcel P. Schilling, Dietrich Kohlheyer, Hanno Scharr, Katharina Nöh, Ralf Mikut

## Abstract

In biotechnology, cell growth is one of the most important properties for the characterization and optimization of microbial cultures. Novel live-cell imaging methods are leading to an ever better understanding of cell cultures and their development. The key to analyzing acquired data is accurate and automated cell segmentation at the single-cell level. Therefore, we present microbeSEG, a user-friendly Python-based cell segmentation tool with a graphical user interface and OMERO data management. microbeSEG utilizes a state-of-the-art deep learning-based segmentation method and can be used for instance segmentation of a wide range of cell morphologies and imaging techniques, e.g., phase contrast or fluorescence microscopy. The main focus of microbeSEG is a comprehensible, easy, efficient, and complete workflow from the creation of training data to the final application of the trained segmentation model. We demonstrate that accurate cell segmentation results can be obtained within 45 minutes of user time. Utilizing public segmentation datasets or pre-labeling further accelerates the microbeSEG workflow. This opens the door for accurate and efficient data analysis of microbial cultures.

## Introduction

Cell-to-cell heterogeneity induced by intrinsic biological factors and extrinsic environmental fluctuations highly affects microbial cell growth and productivity of microbes [1–6]. State-of-the-art microfluidic lab-on-chip systems with highly precise environmental control and time-lapse imaging enable the investigation of causes and consequences of such heterogeneity at the single-cell level [7–9]. Interesting growth parameters include the number of cells over time or size distributions. To extract such quantities from the acquired image sequences, all cells in all images need to be individually segmented, a process that needs to be highly automated and reliable due to the high numbers of cells (up to several tens of thousands of cells per experiment). However, virtually error-free segmentation of acquired time-lapse images remains challenging, as contrast and signal-to-noise ratio are typically low due to limited lighting conditions to avoid phototoxic stress.

Cell segmentation tools are, therefore, often highly specialized to specific imaging conditions. Such specialized tools must be adapted when acquisition settings or experimental parameters change to cope with other cell morphologies or image modalities. When the tools are based on traditional image processing methods, e.g., ChipSeg [10], some adaptations can be achieved by manually adjusting parameters. Unfortunately, this often requires expert knowledge of the underlying method and sometimes even code adaptations. In contrast, deep learning-based methods are not tuned manually and are much more versatile but need to be (re-)trained on annotated data (see Fig 1). Due to this versatility, deep learning methods dominate cell segmentation challenges covering multiple object morphologies and imaging techniques, e.g., the 2018 Data Science Bowl [11]. Furthermore, it is envisioned in [12] that replacing existing analysis pipelines with more accurate deep learning counterparts would greatly aid researchers who conduct live-cell imaging experiments by saving countless person-hours of curation.

**Fig 1.**
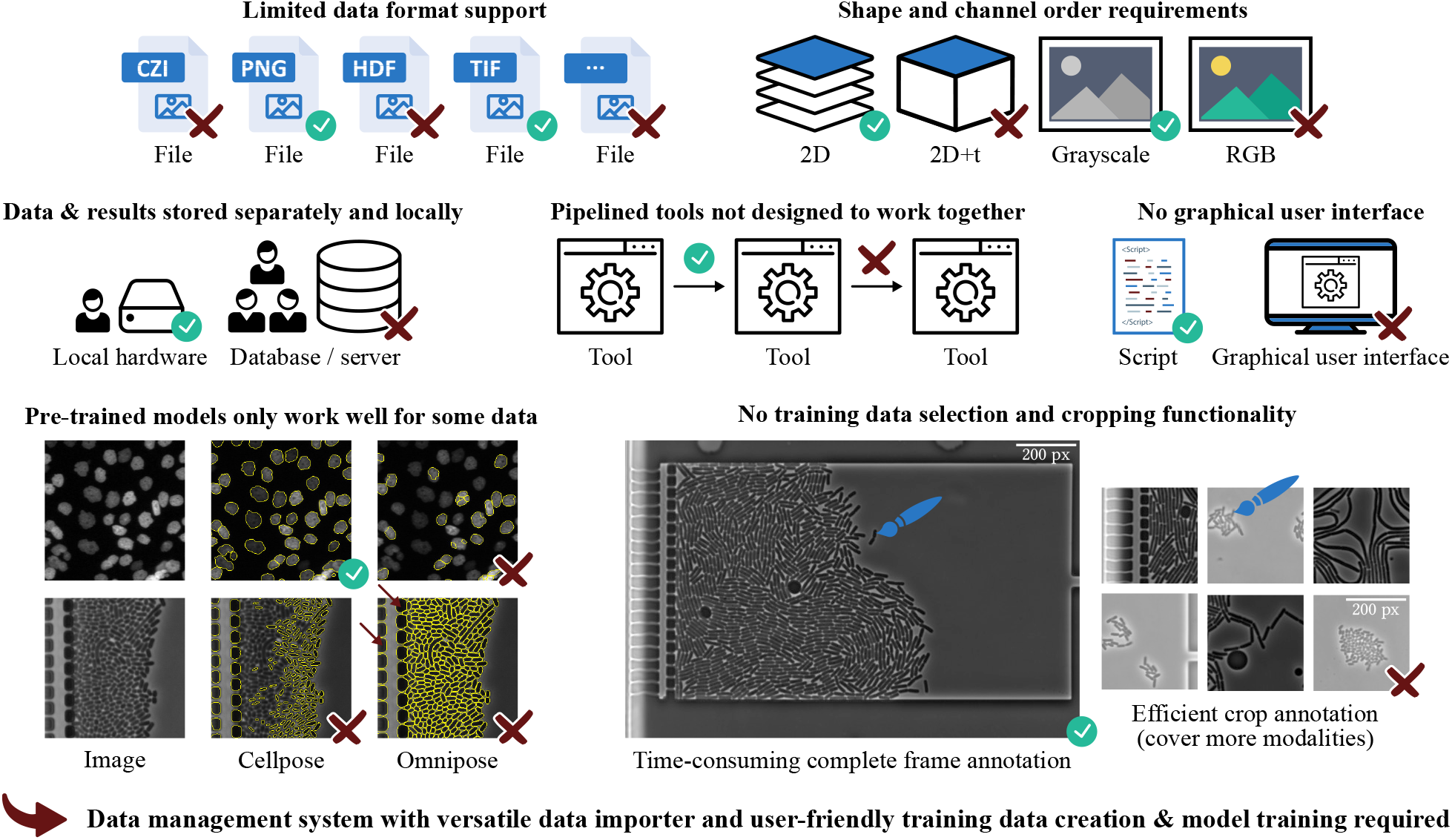
Typical barriers when using segmentation software. Cell segmentation methods are typically designed for specific applications and therefore often lack a data management system with a versatile data importer. This shortcoming results in the need for file format and image shape conversion steps. Furthermore, multiple tools often need to be combined to cover the whole workflow, from training data creation to applying trained models. Again, further processing steps may be required to enable tool compatibility. For many applications, it is not (yet) possible to do without own annotated training data. Interactive cropping functionalities are helpful in this case and enable an efficient annotation for dense-growing organisms. Nevertheless, this feature has not yet been included in cell segmentation software. Note: fluorescence images have been inverted for Omnipose [13], and phase contrast images have been inverted for Cellpose [14] for better results.

Training a deep learning-based method involves a pipeline consisting of multiple steps usually covered by various sequentially applied tools or programming steps. A typical workflow starts with training data creation, data handling and loading, model training, model evaluation – these steps may be iterated until the model is suitably accurate – and finally, the application of trained models to experimental data. Integrating all needed steps in a single, highly automated toolbox is desirable from a user perspective. In particular, the data management, the training data creation, and the training of the deep learning models need to be user-friendly and time efficient, e.g., by using pre-labeling with subsequent manual corrections and full automation of routine steps. Thus far, deep learning approaches for the segmentation of biological data either lack a smooth training data creation and retraining workflow or a data management system like OMERO [15]. Recent examples are Misic for the high-throughput cell segmentation of complex bacterial communities [16] and Omnipose for robust segmentation of bacteria and other elongated cell shapes [14]. Fig 1 summarizes some typical barriers for end users when using cell segmentation software. Moreover, a survey of 704 National Science Foundation principal investigators identified the current and future data analysis needs (i) sufficient data storage, (ii) updated analysis software, (iii) training on data management and metadata, (iv) support for bioinformatics and analysis, and (v) training on basic computing and scripting [17], which shows that there is a need for easy-to-use and state-of-the-art analysis tools with standardized data management.

This paper presents microbeSEG, an open-source deep learning-based segmentation tool that uses the freely available OMERO [15] for data and metadata management. microbeSEG is a tool tailored to instance segmentation of various cell morphologies and imaging techniques. In contrast to other solutions, microbeSEG covers the whole processing pipeline needed for image analysis: users can easily create training datasets, annotate cells with the jointly developed annotation toolkit ObiWan-Microbi [18], and train and apply deep learning models – without any conversion steps or programming. Due to the use of OMERO and its versatile data importer, supporting over 150 image formats, data management is straightforward and user-friendly. Furthermore, we show that accurate cell segmentation is possible even with short training data creation times and without expert knowledge. microbeSEG provides powerful deep learning-based cell segmentation in an easy-to-use tool and is publicly available at https://github.com/hip-satomi/microbeSEG. In the remainder of this work, the microbeSEG functionalities and workflow are introduced, the workflow efficiency is evaluated, and the key features of available cell segmentation software are compared.

## Materials and methods

Fig 2 provides an overview of the microbeSEG functionalities and its needed components, OMERO for data management and ObiWan-Microbi, a jointly developed tool for manual annotation. The key features of our new software are the use of OMERO and the completeness of the workflow, which includes time-efficient interactive training data crop selection, a simple but effective step that has not been included in the software solutions we have reviewed. microbeSEG facilitates deep learning for biologists since no data format conversion steps are needed for data annotation, model training, and inference. Such steps are required when tools need to be combined that are not designed to work together, and no data management system providing file format standards is used. The functionalities and the components of microbeSEG are described in this section.

**Fig 2.**
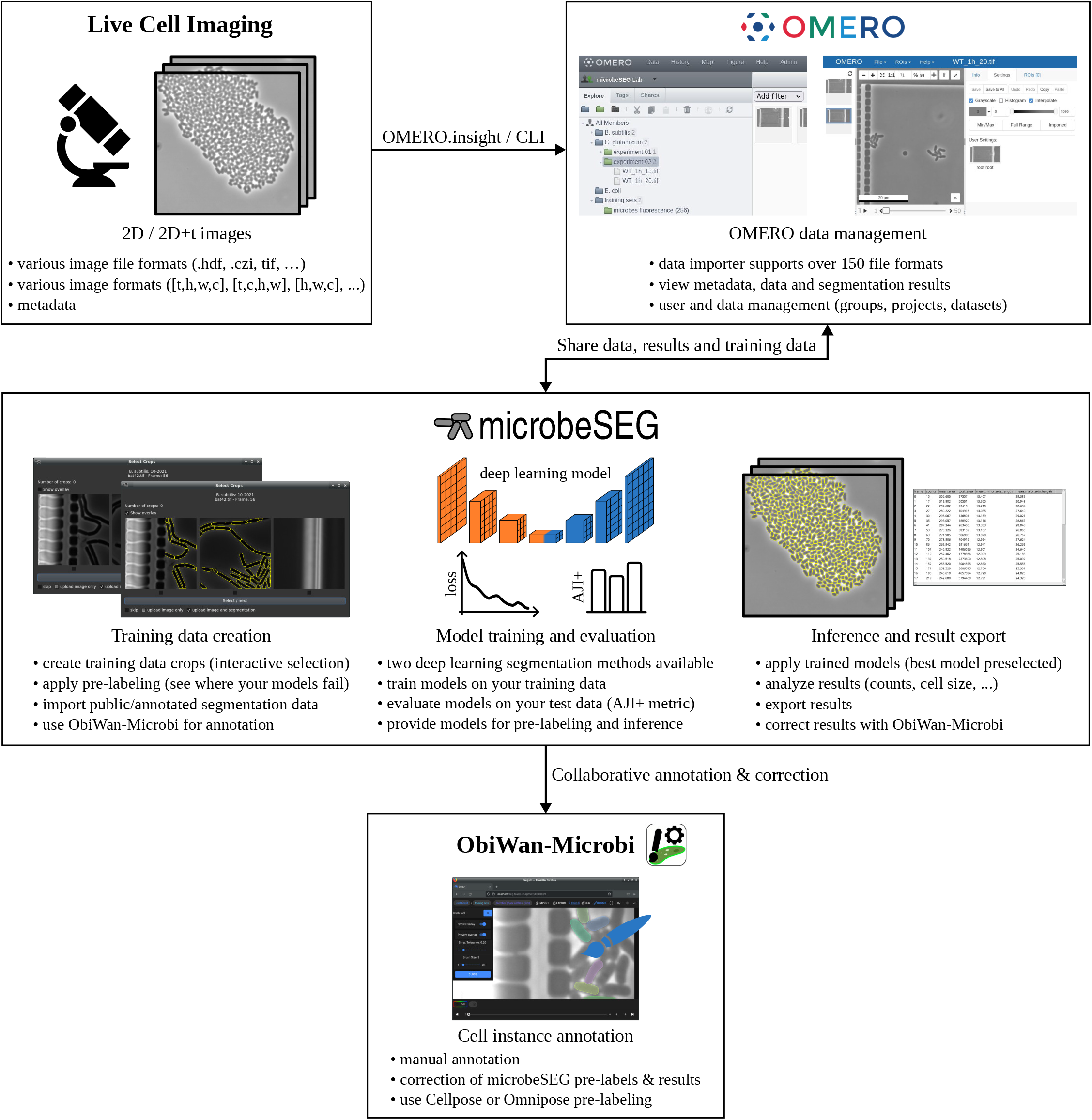
microbeSEG overview. OMERO is used for data management since it provides a versatile data importer and standardizes file handling [15]. Data can be viewed in the browser with the OMERO.web client. microbeSEG offers training data creation, training, evaluation, and inference functionalities. The jointly developed toolkit ObiWan-Microbi is used for manual annotation and result correction [18].

### OMERO data management

OMERO is an open-source software platform for accessing and using a wide range of biological data and provides a unified interface for images, matrices, and tables [15]. Over 150 image formats can be imported with the OMERO.insight desktop client. Thereby, data are organized into projects and datasets. After the import, the data can be processed with microbeSEG. With the OMERO.web client, imported data, microbeSEG training data, and microbeSEG results can easily be accessed and viewed in the browser.

### Training data creation

microbeSEG training sets are managed as OMERO datasets. When adding a new training set to OMERO using the graphical user interface of microbeSEG, a crop size needs to be selected. The crop size selection is required since annotating whole images or frames is tedious and time-consuming, e.g., for frames with microbial colonies of hundreds or thousands of cells. More efficiently is to annotate smaller crops showing fewer cells (see Fig 1) since the cropping enables covering more image and cell features in the same annotation time, resulting in a more diverse training set. Therefore, crop proposals of selected files can be viewed and added to the training set (Fig 3a). Up to three crops – depending on the image and crop size – randomly extracted from non-overlapping image regions are shown per frame and can interactively be selected and uploaded to OMERO. When trained models are already available, pre-labeling allows for identifying areas where the segmentation fails and improvements are needed (Fig 3b). In addition, this approach reduces the annotation time for correctly or partially segmented cells. Selected crops are automatically assigned to a training, a validation or a test subset. However, there is also an option for an explicit assignment. Furthermore, annotated datasets, e.g., publicly available training data, can be imported.

**Fig 3.**
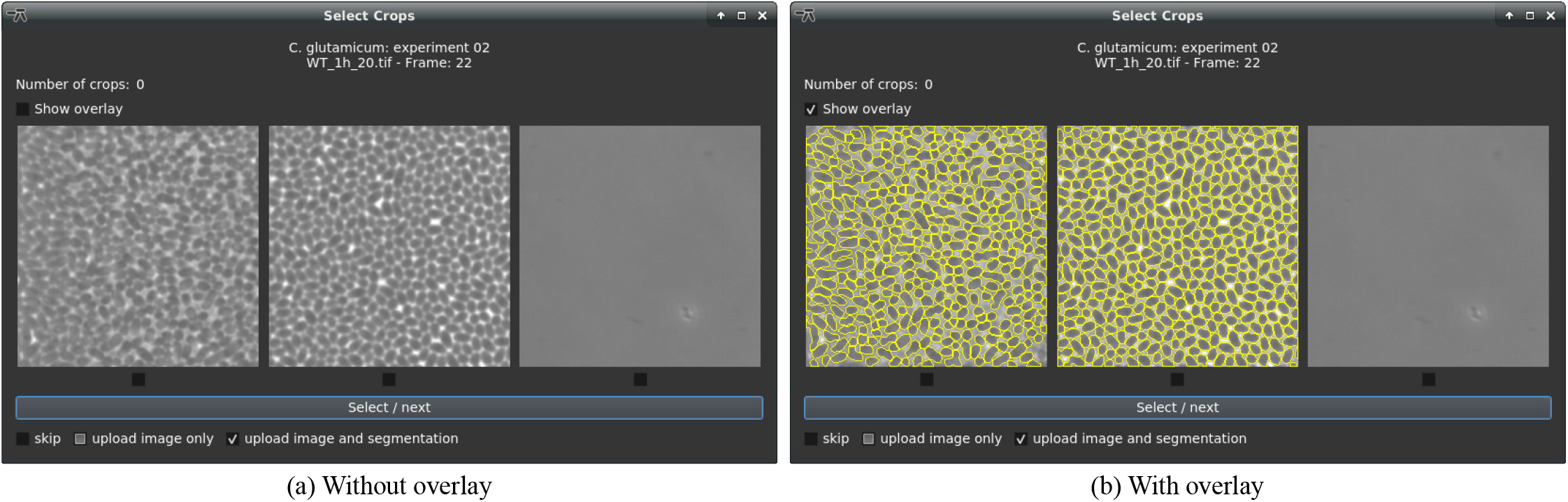
Crop creation interface. Automatically proposed crops can be selected and uploaded to OMERO (a). The crop proposals are extracted randomly from different non-overlapping image regions (the left crop originates from the left image region, and the right crop from the right). For the pre-labeling, it is possible to upload only the image or the image with its prediction (b). The image-only upload is helpful when the pre-label predictions require too many manual corrections.

The training data crops can be annotated with ObiWan-Microbi, an open-source microservice platform for segmentation ground truth annotation in the cloud [18]. The Angular- and Ionic-based web app connects to the OMERO data backend and has been developed jointly with microbeSEG. In addition, to the microbeSEG pre-labeling, Omnipose [14] and Cellpose [13] can be used.

### Model training and evaluation

#### Segmentation methods

In microbeSEG, two deep learning-based instance segmentation methods are implemented: (i) a simple semantic-based method with multi-class output, i.e., background, cell interior, and cell boundary (see Fig 4b), and (ii) a distance transform-based method (see Fig 4c) [19,20]. The latter method (ii) has proven its applicability for phase contrast, bright field, and fluorescence images for diverse cell morphologies in the Cell Tracking Challenge (http://celltrackingchallenge.net/) [19, 21] where the method currently holds nine top-three rankings as participants KIT-GE (2) and KIT-GE (3). The simple boundary method is easy to interpret and serves as a baseline. Both methods are based on the U-Net convolutional neural network [22], with the distance method using two decoders, one for each predicted output. The single-decoder U-Net has about 34 million parameters and the double-decoder U-Net has 46 million. However, the network parameters are automatically reduced to a minimum of 2 million and 3 million, respectively, if not enough memory is available. In the seeded watershed-transform-based post-processing, touching instances are split using the cell boundary prediction or the neighbor distance prediction. The distance method post-processing has two parameters: a threshold for adjusting the cell size and a threshold for adjusting the seed extraction.

**Fig 4.**
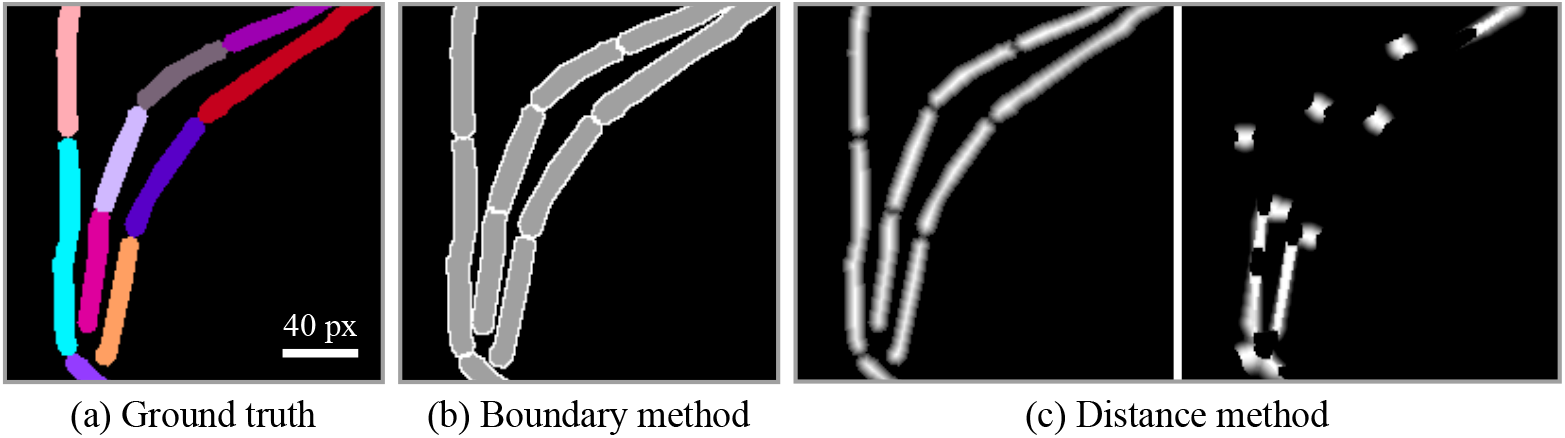
Training data representations of the two microbeSEG methods. From the ground truth (a – instances: color-coded), the boundary representation (b – cell interior: gray, cell boundary: white) and the distance representations (c – distance to background, inverse distance to neighbors) can be computed. A deep learning model is trained to predict either the boundary representation or the distance representations, and the single instances are recovered in the post-processing.

#### Training

For training on a selected annotated training set, the segmentation method, the batch size, the optimizer, and how many models should be trained need to be specified. Adam [23] and Ranger [24] are available as optimizers, each with its own pre-defined settings, i.e., start learning rate, minimum learning rate, and learning rate scheduler (reduce learning rate on validation loss plateau). The default settings are: distance method, Ranger optimizer, training of five models, and a batch size of four. Usually, the default settings are sufficient, but the user may reduce the batch size if memory availability is limited. Flip, rotation, scaling, contrast, blur, and noise augmentations are applied during training. The maximum number of training epochs is set automatically depending on the dataset and crop size. In addition, a stopping criterion depending on the maximum number of training epochs and validation loss improvement is applied. A weighted sum of cross entropy loss and channel-wise Dice loss is used for the boundary method. The distance method is trained with a smooth *L*_1_ loss.

#### Evaluation

Trained models can be evaluated on the automatically split test set. Thereby, appropriate parameters are set for the post-processing of the distance method, i.e., the cell size adjustment and the seed extraction threshold. A slightly modified version of the aggregated Jaccard index AJI [25] is used as the evaluation measure. The modification AJI+ prevents overpenalization by using a one-to-one mapping instead of a one-to-many mapping for the predicted objects [26, 27]. The AJI+ score ranges from 0 to 1, with 1 indicating perfect segmentation. The mean AJI+ score and the standard deviation over the single test images of each evaluated model are saved in a csv file.

### Inference and result export

The best evaluated model with its parameter set is selected automatically for inference. However, the user can also select the model manually. If a not yet evaluated model is selected, the default thresholds are used for the distance method, which may result in (slightly) too large or too small cells for this method. The segmentation results can be attached to the corresponding OMERO image as polygon regions of interest, which enables joint storage of images and results, and can be viewed with the OMERO.web client. In addition, the cell counts, mean cell area, mean cell minor and major axis length, and the total cell area are determined. The result export includes: the original image (.tif), intensity-coded instance segmentation masks (.tif), a cell outlines image (.tif), the original image overlaid with the cell outlines (.tif), and analysis results (.csv).

### Implementation, installation, and dependencies

microbeSEG is implemented in Python [28] and uses PyTorch as the deep learning framework [28,29]. The graphical user interface is built with PyQt [30]. The software and the source code are available under the MIT license in our code repository. A requirements file and a step-by-step guide are provided in the code repository for easy installation and usage. Access to an OMERO server is required for running microbeSEG. For testing purposes, a demo server account can be requested from the OME team in Dundee: http://qa.openmicroscopy.org.uk/registry/demo_account/. Furthermore, an OMERO server can be set up during the ObiWan-Microbi installation, which is available at https://github.com/hip-satomi/ObiWan-Microbi.

### microbeSEG dataset

Microbe data shown in this paper were acquired with a fully automated time-lapse phase contrast microscope setup using a 100x oil immersion objective. Cultivation took place inside a special microfluidic cultivation device [31]. The assembled microbe training, validation, and test sets, including 2930 cells, are available online at https://doi.org/10.5281/zenodo.6497715 (see next section).

## Results

### microbeSEG workflow efficiency

Table 1 shows the microbeSEG segmentation accuracy in terms of the aggregated Jaccard index AJI+ in dependence of the annotation time on a test set consisting of two microbe types, i.e., of 721 *B. subtilis* and 1168 *E. coli* cells. The 24 test images of size 320 px × 320 px, 12 for each species, were annotated by three experts, who cross-checked their annotations. The microbeSEG user selected and annotated crops from different experimental data than the experts used. The distance method parameters have been adjusted automatically on the internal test set of the microbeSEG user. Each of the ten boundary models ended training within 6 minutes, and each of the ten distance models within 18 minutes on a system with an Nvidia Titan RTX (batch size: four).

**Table 1.**
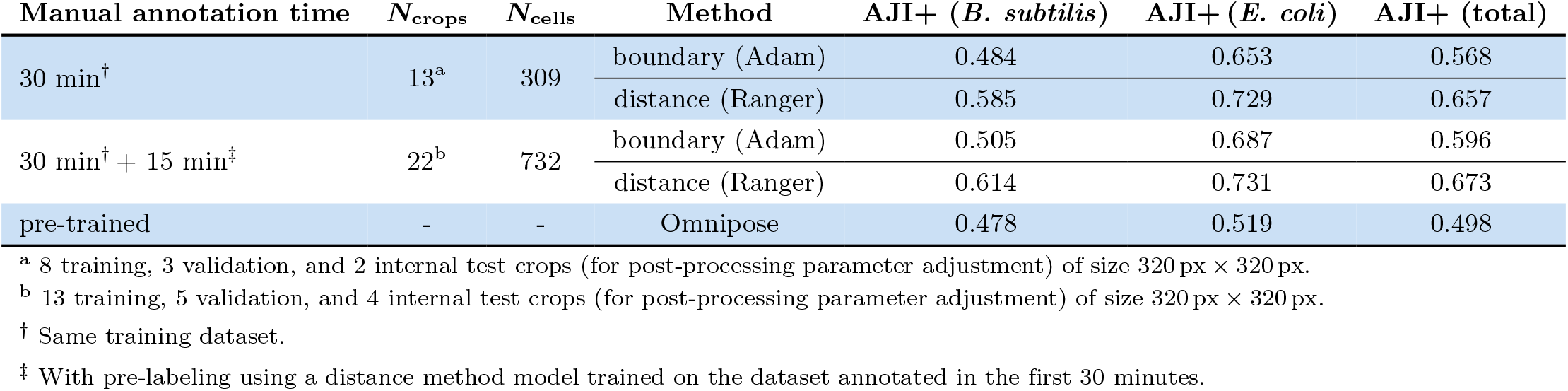
microbeSEG accuracy in dependence of annotation time. All methods are evaluated on 12 *B. subtilis* and 12 *E. coli* test images. For each microbeSEG setting, the median aggregated Jaccard index AJI+ out of five trained models is shown. The times include the crop selection.

Switching from completely manual annotation to correcting pre-labeled image crops after the initial annotation phase results in more cells annotated per time. Therefore, a model trained on the dataset annotated in the first 30 minutes has been used. The additional training data further increase the scores for both cell types, especially for the simpler boundary method, which requires larger training datasets. The default microbeSEG settings – distance method and Ranger optimizer – yield better segmentations than the boundary method. Fig 5 shows exemplary segmentations for the default settings, indicating that reasonable segmentation results for the two microbe types can be obtained within 30 minutes of annotation time, even without utilizing public datasets, pre-training, or pre-labeling. For comparison, Omnipose [14], a deep learning method pre-trained on a large database of bacterial images with 27 000 annotated cells, yields convincing segmentations of cells but, without retraining, suffers from the detection of background structures.

**Fig 5.**
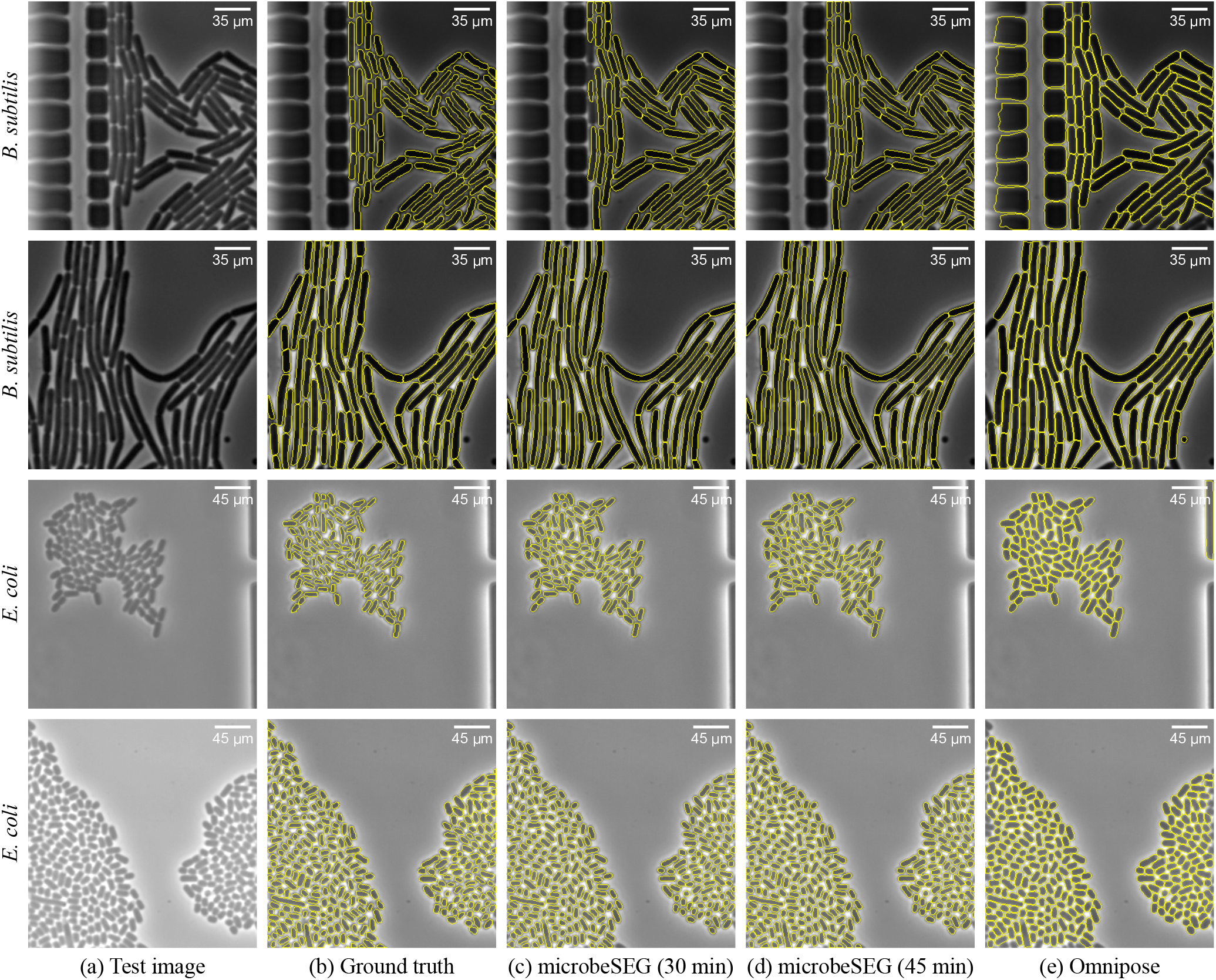
Exemplary images and segmentation overlays for *B. subtilis* and *E.coli*. Shown are the results of the median distance method microbeSEG models from Table 1.

### Qualitative U2OS cell segmentation results using imported HeLa cell training data

The import of annotated datasets helps to decrease manual interaction time further or even makes the need for manual annotations or corrections obsolete in some cases where very similar training data is available. Fig 6 shows qualitative segmentation results of U2OS cells [32] with a microbeSEG model trained on HeLa cell data from the Cell Tracking Challenge [33, 34]. Even without annotated U2OS cells, a good segmentation quality is achieved.

**Fig 6.**
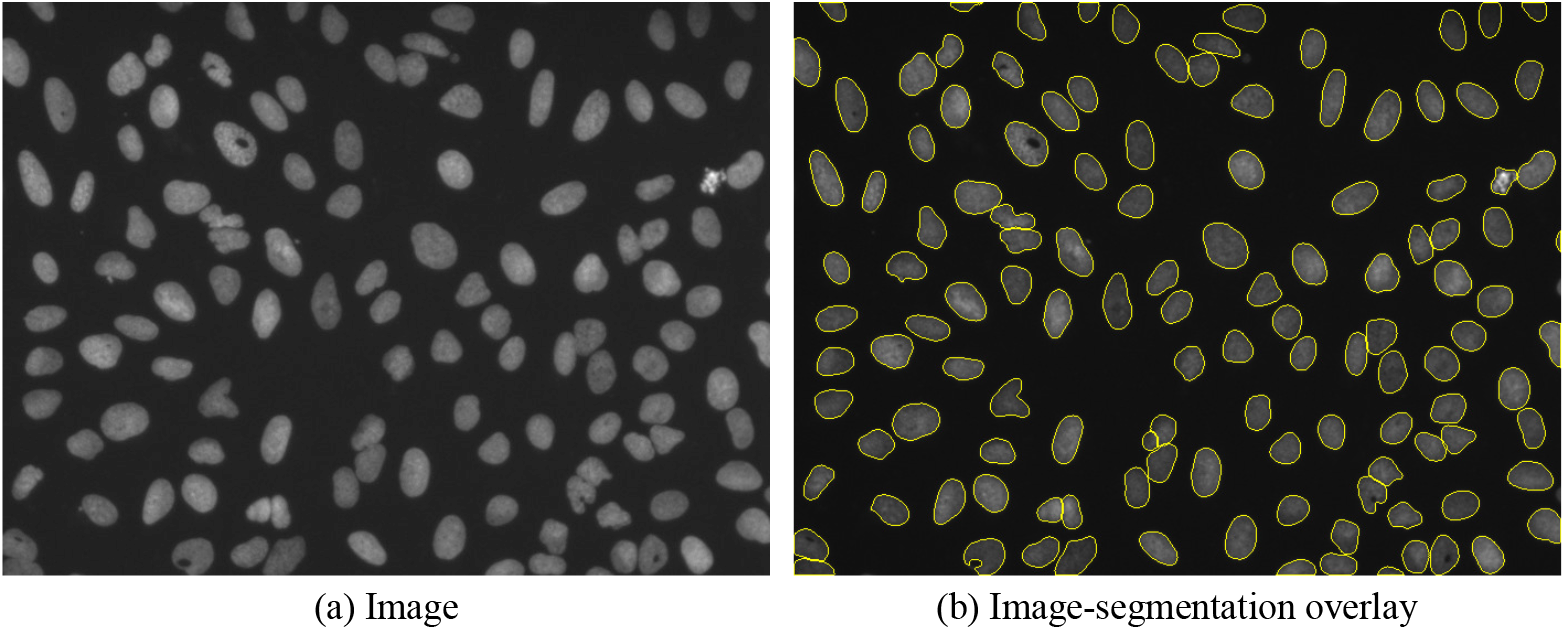
Segmentation of U2OS cells from the BBBC039 dataset [32]. The microbeSEG segmentation model (default settings) has been trained on HeLa cells from the Cell Tracking Challenge [33, 34].

### Data analysis

For time-lapse experiments, basic analysis results, i.e., cell counts, mean cell area, mean cell minor/major axis length, and total area, can be exported after segmentation. Possibly needed manual corrections to reach a virtually error-free segmentation can be applied using ObiWan-Microbi. The results can easily be plotted using Fiji [35], as Fig 7 shows.

**Fig 7.**
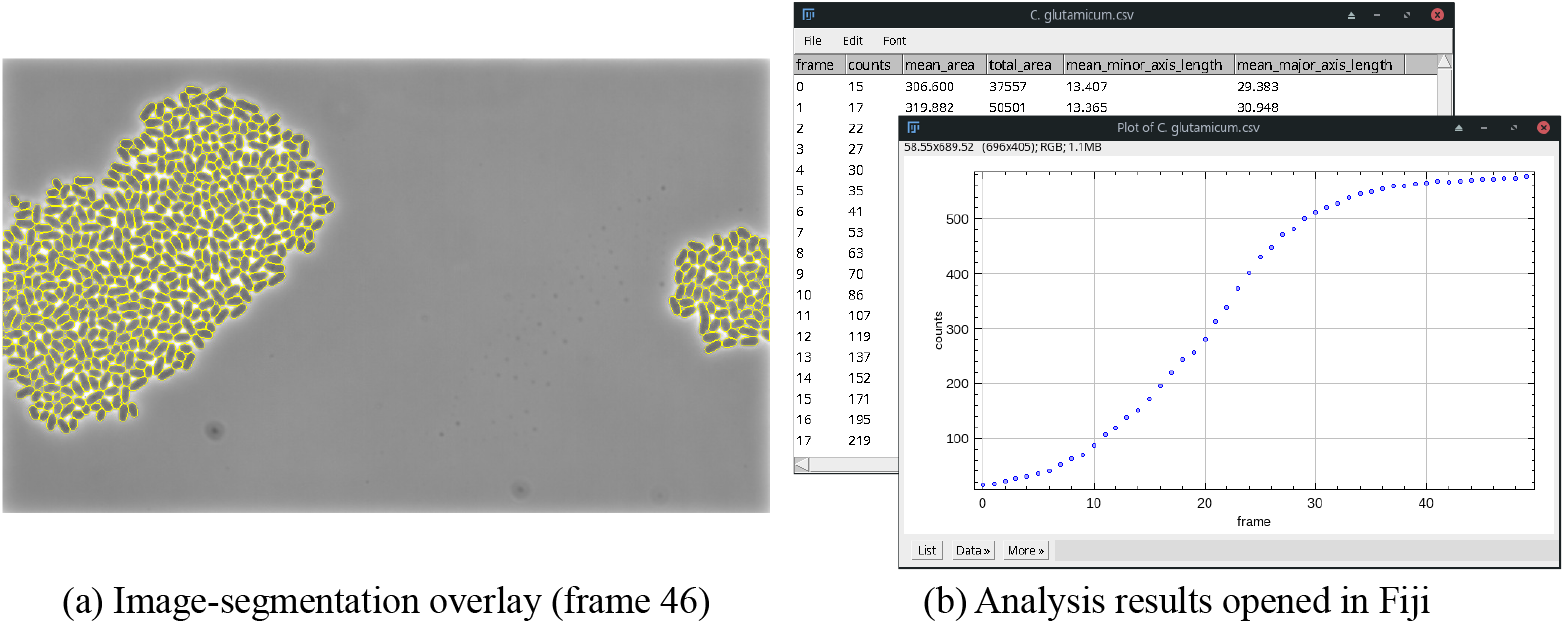
Exemplary microbeSEG segmentation (a) and analysis results (b) for a growing *C. glutamicum* colony. The results can easily be viewed in Fiji.

### Segmentation tools feature comparison

Table 2 provides an overview of crucial features we believe a cell segmentation tool should have for ease of use and also shows which software meets these features. Some tools support only a limited number of data formats, which results in additional conversion steps before the software can be used. In addition, most tools use no data management system that can store, visualize, and share data, metadata, and results. However, the support of such a data management system and its corresponding file format standards facilitates the use of segmentation tools. So far, most deep learning software with a graphical user interface focuses on applying pre-trained models, and model training is not directly possible and requires programming expertise. Furthermore, training data annotation can require the use of incompatible annotation tools leading to further conversion steps. microbeSEG is the only tool that covers all key features (together with the jointly developed ObiWan-Microbi).

**Table 2.** Cell segmentation software key feature comparison. Considered are only tools with a graphical user interface since end users should not need programming expertise. Non deep learning segmentation methods may require expert knowledge for parametrization and are not state-of-the-art anymore. Data format support does not necessarily mean that each image can be processed: if no data management system (DMS) with metadata support is used, e.g., the channel dimension can be the first or the last dimension for .tif files, and the method may have requirements on the channel dimension position. **−**: feature not fulfilled/supported, 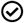: feature only fulfilled/supported with restrictions, **✓**: feature fulfilled/supported.

| Software | DMS | Data formats | Deep learning | Cropping | Annotation | Pre-labeling | Training | Result corr. |
| --- | --- | --- | --- | --- | --- | --- | --- | --- |
| AutoCellSeg [36] | — | .jpg, .png, .tif, .bmp | — | — | — | — | — | ✓ |
| BacStalk [37] | — | .jpg, .tif | — | — | — | — | — | —<sup>a</sup> |
| Cellpose/Omnipose [13, 14, 38] | — | .gif, .jpg, .png, .tif | ✓ | — | ✓ | ✓ | ✓ | ✓ |
| ChipSeg [10] | — | .tif | — | — | — | — | — | — |
| DeepImageJ [39] (Fiji plugin [35]) | — | > 150<sup>b</sup> | ✓ | — | — | — | — | — |
| Misic [16] (napari plugin [40]) | — | > 150<sup>b</sup> | ✓ | — | ☑<sup>c</sup> | — | — | — |
| Orbit [41] | ✓<sup>d</sup> | > 150<sup>b</sup> | ☑<sup>e</sup> | — | ✓ | — | ☑<sup>f</sup> | — |
| StarDist [42] (napari plugin [40]) | — | > 150<sup>b</sup> | ✓ | — | ☑<sup>c</sup> | — | ☑<sup>g</sup> | — |
| YeaZ [43] | — | 25 | ✓ | — | — | — | — | ✓ |
| YeastSpotter [44] | — | N.A. | ✓ | — | — | — | — | — |
| microbeSEG | ✓<sup>d</sup> | > 150<sup>b</sup> | ✓ | ✓ | ✓<sup>h</sup> | ✓ | ✓ | ✓<sup>h</sup> |

<sup>a</sup> Only deletion of objects, no shape refinement or correction of false negatives, merges, and split objects.
<sup>b</sup> OME Bio-Formats support (napari needs the napari-aicsimageio plugin: https://www.napari-hub.org/plugins/napari-aicsimageio).
<sup>c</sup> In principle possible with napari (plugins) but no training functionality with the graphical user interface.
<sup>d</sup> OMERO.
<sup>e</sup> With copy and paste of a segmentation script into the script editor: https://www.orbit.bio/deep-learning-object-segmentation/
<sup>f</sup> Only with a Python script outside of Orbit.
<sup>g</sup> Not within the graphical user interface, but in principle within the napari IPython console.
<sup>h</sup> With the associated tool ObiWan-Microbi.

## Discussion

Utilizing ObiWan-Microbi for training data annotation, microbeSEG covers the whole pipeline from training data creation, training, evaluation, and inference. Both tools are designed to work together seamlessly and use OMERO for data management. The optimized workflow allows producing high-quality segmentation results in reasonable annotation time. Our results indicate that switching to pre-label corrections is a better strategy than annotating manually when suitably accurate models have been trained since more cells can be annotated per time. In our experiments, we observed that it is a good strategy to correct mainly object-level errors, i.e., merges, splits, and added or missing cells. As the qualitative segmentation results show, the AJI+ scores for *B. subtilis* are mainly limited by size differences and different handling of cell division events. This discrepancy is due to the different decision boundaries of the microbeSEG user and the individual expert annotators in this experiment, especially for the cell division events of the challenging filamentous *B. subtilis* cells and for the *E. coli* size. We assume that when the same person annotates training and test data, the AJI+ scores rise significantly.

The Omnipose segmentation results show that available software solutions may need retraining even when pre-trained on a large dataset, e.g., to avoid the segmentation of background structures on a microfluidic device. However, such solutions can further reduce the manual annotation time. For instance, ObiWan-Microbi includes Omnipose and Cellpose pre-labeling. The prerequisite for saving annotation time is that these solutions work quite well for the present data. A drawback is, however, that object shapes may not match as well as with manual annotation or the microbeSEG pre-labeling, which enables learning the annotator’s object size decision boundary.

Once a large and diverse annotated dataset is established, further annotation and model training is required only for significant changes in the experimental setup or for specific cases where the segmentation fails. Thereby, the microbeSEG crop selection with pre-labeling allows adding exactly those new training data where segmentation errors occur. A drawback of the training data crop creation is that for large image sizes, a small selected crop size, and low object density, all proposed crops of a frame could show no cells. This behavior can for example occur in a 2048 px × 2048 px image with the smallest selectable crop size of 128 px × 128 px × and less than ten small objects. In such a case, a larger crop size (crop sizes up to 1024 px × 1024 px are available) needs to be selected. Another limitation of microbeSEG is that the segmentation method does not, like many others, support the segmentation of overlapping objects.

In summary, we believe that the major concern about deep learning for cell segmentation, which is that extensive and time-consuming annotation of training data is required, has been overcome with our tool microbeSEG, which provides an efficient workflow and an accurate segmentation method. Our segmentation software feature comparison has shown that no other segmentation tool provides all functionalities needed for the efficient segmentation of microbes. Certainly, microbeSEG can also be applied for the segmentation of cell nuclei or other objects. To further facilitate the data annotation for microbeSEG users, label inspection strategies [45] and training data simulators are of particular interest. A long-term goal is to leverage deep learning for 3D applications, e.g., to study plant-microbe interactions [46]. Integrating other state-of-the-art segmentation methods like Omnipose or StarDist into our tool with its complete workflow is another possible future direction.

## Acknowledgment

We are grateful for the funding by the Helmholtz Association in the programs Natural, Artificial and Cognitive Information Processing (T.S., R.M.), Engineering Digital Futures: Supercomputing, Data Management and Information Security for Knowledge and Action (H.S.), HIDSS4Health - the Helmholtz Information & Data Science School for Health (R.M.), and the Helmholtz Association Initiative and Networking Funds through Helmholtz AI (O.N., R.M.) as well as the Helmholtz Imaging Platform within the project SATOMI (J.S., H.S., D.K., K.N., R.M.). Funded by the Deutsche Forschungsgemeinschaft (DFG, German Research Foundation) – 491111487 (K.N.). The funders had no role in study design, data collection and analysis, decision to publish, or preparation of the manuscript.

## Notes

### Competing Interest Statement

The authors have declared no competing interest.

### Summary of Updates

Figures revised (readability/clarity); information about model parameters added; materials and methods section restructered; introduction expanded.

